# Improved immunoassay sensitivity and specificity using single-molecule colocalization

**DOI:** 10.1101/2021.12.24.474141

**Authors:** Amani A. Hariri, Sharon S. Newman, Steven Tan, Dan Mamerow, Michael Eisenstein, Alexander Dunn, H. Tom Soh

## Abstract

Enzyme-linked immunosorbent assays (ELISAs) are a cornerstone of modern molecular detection, but the technique still suffers some notable challenges. One of the biggest problems is discriminating true signal generated by target molecules versus non-specific background arising from the interaction of detection antibodies with the assay substrate or interferents in the sample matrix. **Si**ngle-**M**olecule **C**olocalization **A**ssay (SiMCA) overcomes this problem by employing total internal reflection fluorescence (TIRF) microscopy to quantify target proteins based on the colocalization of fluorescent signal from orthogonally labeled capture and detection antibodies. By specifically counting colocalized fluorescent signals, we can essentially eliminate the confounding effects of background produced by non-specific binding of detection antibodies. We further employed a normalization strategy to account for the heterogeneous distribution of the capture antibodies, greatly improving the reproducibility of our measurements. In a series of experiments with TNF-α, we show that SiMCA can achieve a three-fold lower limit of detection compared to conventional single-color assays using the same antibodies and exhibits consistent performance for assays performed in complex specimens such as chicken serum and human blood. Our results help define the pernicious effects of non-specific background in immunoassays and demonstrate the diagnostic gains that can be achieved by eliminating those effects.

## Introduction

Even after nearly 50 years, the enzyme-linked immunosorbent assay (ELISA) remains an essential tool for detection of protein biomarkers for both basic research and clinical diagnostics^1,2^. The ELISA has changed relatively little since its inception in 1971—two different antibodies are used to capture and label the target in a “sandwich” format that generates a signal only when both the capture antibody (cAb) and detection antibody (dAb) are bound to the target^1^. This dual-binding requirement confers excellent specificity, but ELISAs remain vulnerable to background signal arising from non-specific binding of antibodies or interferent proteins to the assay substrate^2^. Unwanted background binding can be mitigated to some extent through the use of more stringent wash conditions, blocking of exposed assay substrate surfaces, or careful management of the amount of dAb^3^. However, these solutions entail trade-offs in terms of assay performance. For example, overly stringent washing can undermine assay sensitivity by causing loss of signal, while the use of insufficient dAb concentrations will undermine the assay’s ability to accurately resolve target concentrations^4, 5^.

Consequently, ELISA-based molecular detection is limited by the challenge of discriminating true target-binding events from non-specific background. Researchers have devised a number of different strategies to overcome this difficult problem.^6^ For example, Chaterjee *et al*. developed a ‘kinetic fingerprinting’ assay for the detection of individual surface-immobilized protein molecules.^7^ Their approach combines a conventional cAb with a Fab antibody fragment probe for detection.^8^ *In vitro* selection is used to isolate probes that exhibit sufficiently fast dissociation kinetics to achieve rapid, repetitive binding to their target, enabling discrimination of true target recognition events from non-specific background. This technique offers exceptional sensitivity in serum, but only a limited number of probes are currently available and the process of generating novel Fab probes remains challenging and resource-intensive. Zhang *et al*. reported the use of aptamers for the detection of small-molecule analytes^9^. By splitting an aptamer that binds to ATP and labeling each fragment with differently colored fluorophores, they were able to develop an assay that reports binding only when the two fluorophores are in proximity to each other, and thus eliminates background from non-specific binding events. This method is limited by the need for split-aptamer probes that can bind to a target analyte with high affinity and specificity. These reagents remain challenging to engineer, and only a small number of such probes have been described in the literature. Furthermore, this approach has only been applied to small-molecule detection, and it remains unclear how well such an assay would perform with protein targets.

In this work, we describe a two-color sandwich immunoassay that discriminates between specific and non-specific binding and can be applied to a wide range of protein analytes. **Si**ngle-**M**olecule **C**olocalization **A**ssay (SiMCA) employs cAbs and dAbs that have been labeled with distinct fluorophores. The sample is imaged with total internal reflection fluorescence (TIRF) microscopy at sufficiently low concentrations of cAb and dAb such that single molecules of each species can be readily imaged (**Fig. 1A**). By discarding dAb molecules that are not colocalized with a cAb counterpart, we can greatly decrease the background signal due to non-specific binding, resulting in an improved signal-to-noise ratio, decreased limit of detection (LOD), and increased accuracy for analyte calibration curves. In addition to non-specific dAb binding, heterogeneous cAb surface loading can also contribute substantially to assay variability. Single-molecule imaging allowed us to normalize dAb counts to the counts of cAb for every field of view, thus overcoming this heterogeneity problem. This approach results in far greater sensitivity and consistency of signal across experiments, even in environments with high background. For example, we demonstrated SiMCA with a pair of well-characterized tumor necrosis factor α (TNF-α) antibodies and showed that we could achieve a three-fold lower LOD in serum relative to a non-colocalization-based assay using the same antibodies (7.6 ± 1.9 pM versus 26 ± 5.8 pM). Furthermore, our measurements remained consistent whether the assay was performed in buffer, 70% chicken serum, or 70% whole human blood. Collectively, these results demonstrate that SiMCA can overcome the sensitivity and reproducibility limitations imposed by non-specific binding, enabling accurate detection of picomolar concentrations of protein even in highly complex biological matrices.

**Figure 1:**
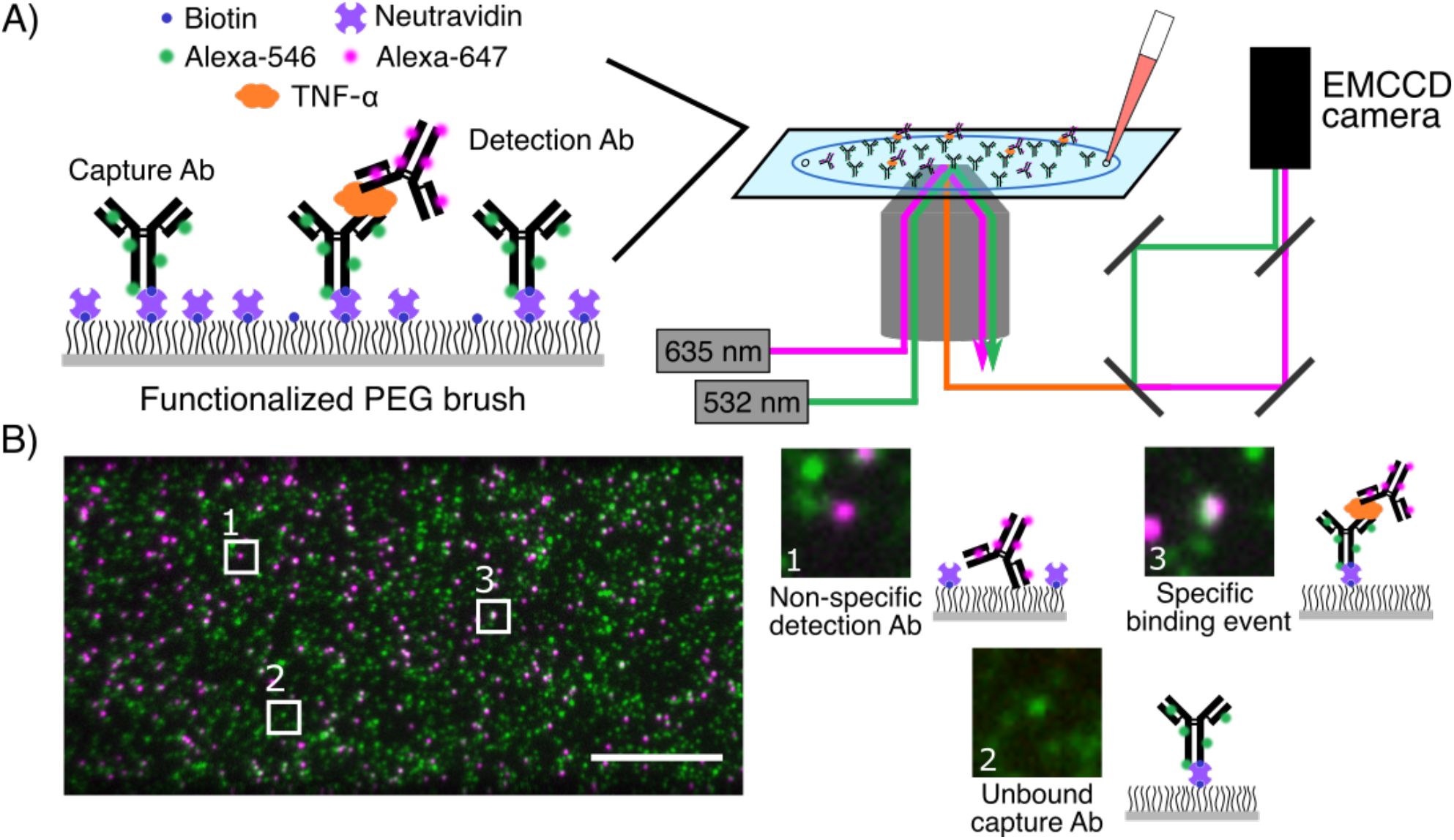
SiMCA platform design. **A**) A glass coverslip is passivated with a mixture of PEG and PEG-biotin, and then treated with neutravidin and biotinylated, Alexa-546-tagged capture antibodies (cAbs). The surface is then incubated with a solution of the target biomolecule and Alexa-647-labeled detection antibody (dAb). The coverslip is imaged using two-color TIRF microscopy. **B)** Images of the cAb and dAb channels are acquired, registered, and analyzed to discriminate non-specific dAb binding (1) and unbound cAbs (2) from true binding events (3). Scale bar = 10 µm.

## Results and Discussion

### Overview of SiMCA

SiMCA is a sandwich-based assay in which cAbs and dAbs are each conjugated to a distinct fluorophore tag, and true binding events are indicated only when both fluorescent signals are colocalized (**Fig. 1A**). As a demonstration, we employed a pair of antibodies that specifically recognize the inflammatory cytokine TNF-α. We labeled the cAb with a green fluorophore (Alexa-546), and site-specifically tagged the antibody with biotin so that it can be immobilized onto a neutravidin-coated surface while ensuring that the antigen-binding domain is appropriately oriented for target binding. The dAb was labeled with a red fluorophore (Alexa-647) (see **Methods, SI Fig. S1)**.

For the assay substrate, we passivated a coverslip with a mixture of PEG and PEG-biotin^10^ to minimize non-specific binding events and to specifically immobilize the biotinylated cAbs via neutravidin-biotin binding. We then incubated the coverslip with a mixture of TNF-α and the dAb and employed a custom two-color TIRF microscope to acquire images via sequential excitation of the green and red dyes with 532- and 635-nm lasers, respectively. Unbound cAbs are detected solely in the green channel, while dAbs that are non-specifically bound to the substrate are registered only in the red channel (**Fig. 1B**). In contrast, true binding events give rise to a ternary sandwich complex, resulting in colocalized red and green signals. We used an automated method for image segmentation and registration to count the single-color dAb signals and colocalized binding events in a high-throughput manner across many fields of view per coverslip.

### SiMCA mitigates non-specific binding and improves reproducibility

It is well known that the use of excess concentrations of dAb in an ELISA can contribute greatly to non-specific binding^4, 11^. Decreasing the level of dAb employed can mitigate this problem, but at the cost of reduced sensitivity due to loss of signal. SiMCA provides the opportunity to measure the extent of non-specific binding and its impact in terms of background signal, as well as the means to remedy that problem. We first assessed the extent of non-specific binding that occurs at various concentrations of dAb in the absence of TNF-α. We incubated cAb-coated coverslips overnight with either low (50 nM) or high (500 nM) concentrations of dAb, washed the coverslips to remove any unbound dAb, and then acquired 128 51.2 µm x 25.6 µm fields of view (FOVs) for each coverslip.

A randomly selected small section of a single FOV shows minimal dAb recruited to the coverslip at 50 nM dAb (**Fig. 2A**, left). In contrast, at 500 nM dAb, we saw a substantial increase in dAb counts. As there was no TNF-α present, these counts solely represent non-specific binding events. As expected, when we overlaid the two fluorescence channels, most dAb spots did not colocalize with a cAb (**Fig. 2A**, middle). For visualization, we inverted the color scale, such that overlapping cAb and dAb spots appear as black spots (**Fig. 2A**, bottom). Any dAb spots that were not spatially overlaid with a cAb are non-specific binding events that would, with conventional methods, be mistakenly counted as a binding event **(SI Fig. S2-3)**. Quantifying the total number of dAbs measured versus the number that colocalized with a cAb across all 128 FOVs of each coverslip revealed that colocalization could eliminate virtually all of these spurious binding events (**Fig. 2B**; **SI Fig S4**). Measured using a single-color, mean dAb counts increased from 0.4 ± 0.6 molecules per FOV at 50 nM dAb to 92 ± 23 molecules at 500 nM dAb. In contrast, colocalized dAb and cAb counts were essentially unchanged, increasing only slightly from 0 ± 0.0 molecules at low [dAb] to 2 ± 1.3 molecules at high [dAb]. These results demonstrate that the SiMCA two-color localization strategy can greatly mitigate the effects of non-specific binding.

**Figure 2:**
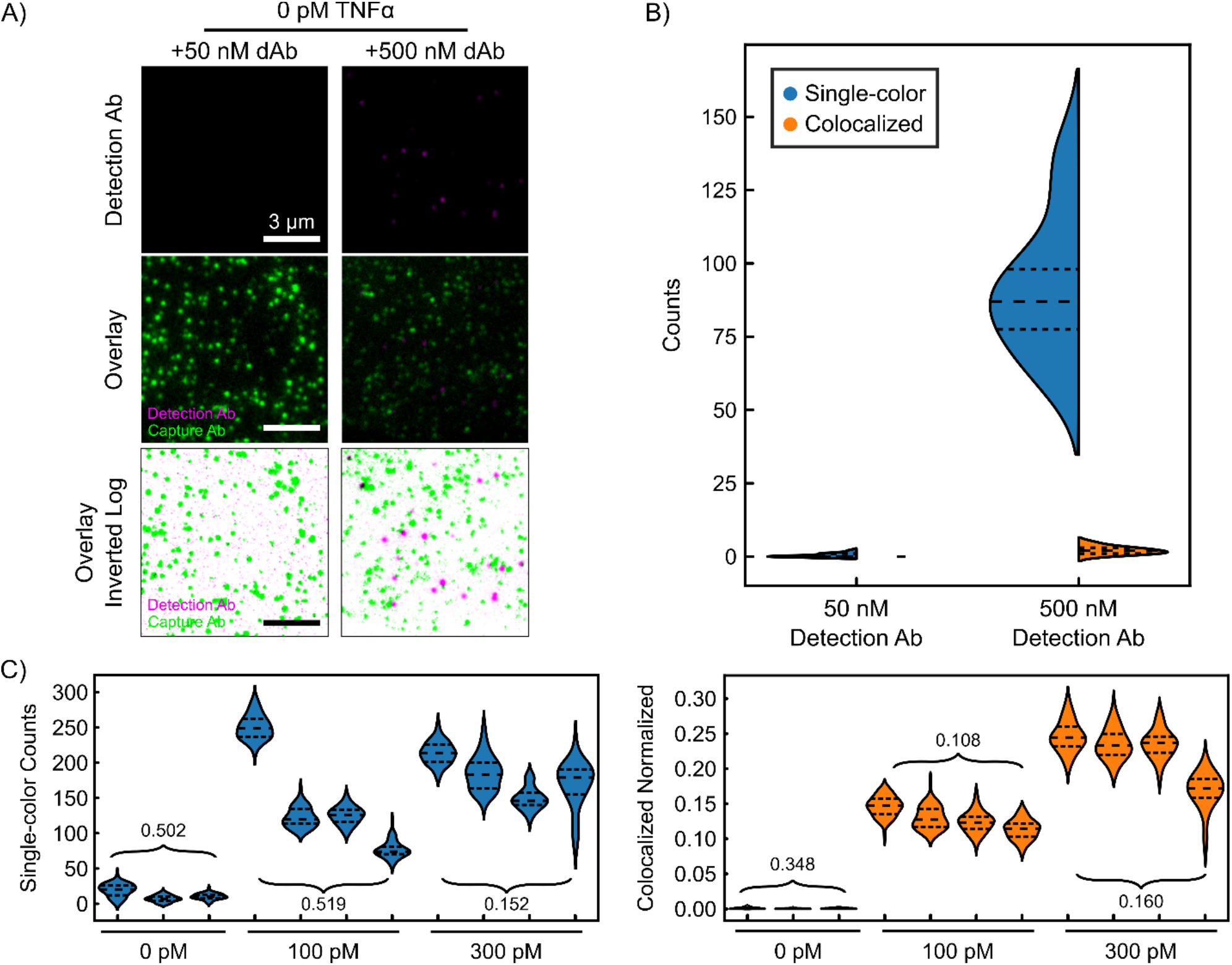
Colocalization minimizes background from excess detection antibody. **A**) Single-color fluorescence images of dAb only (*top*), two-color images of cAb and dAb (*middle*), and log-scale inverted composite images of two-color detection (*bottom*) in the absence of TNF-α and with 50 nM (*left*) or 500 nM (*right*) dAb. Dark spots in the bottom panels represent colocalized signal from the two fluorophores. **B)** Distributions of absolute single-color and colocalized counts across 128 fields of view (FOVs). Dashed lines demarcate quartiles of the distribution. **C)** Absolute number of dAbs per fields of view (*left*) and normalized, colocalized counts (*right*) across different coverslips and TNF-α concentrations. Each violin represents 128 FOVs per coverslip. Numbers are the coefficients of variance (CV) of the mean FOV counts across coverslips.

The vulnerability of single-color methods to non-specific background is further exacerbated by coverslip heterogeneity, wherein the stochastic distribution of cAb on the surface may lead to discrepancies in the number of colocalized pairs observed with identically prepared coverslips and samples. To explore this effect, we incubated multiple coverslips functionalized with cAb with varying levels of TNF-α (0, 100, and 300 pM) and 50 nM dAb. Quantifying the absolute numbers of dAb counts per field of view across coverslips resulted in a high coefficient of variance (CV) at 100 pM TNF-α (**Fig. 2C**, left), whereas counting only colocalized events resulted in slightly lower CVs due to reduced background (**SI Fig. S5)**. This modest reduction is attributable to the fact that cAb counts varied considerably both across multiple FOVs within a single coverslip as well as across coverslips (**SI Fig. S6)**. To account for this heterogeneity, we normalized the colocalized dAb counts by the cAb counts in each FOV, where a normalized count of 1 is the theoretical maximum binding and 0 is the theoretical minimum. This normalization greatly decreased the signal variance, with a notable 4.8-fold reduction in CV for the 100 pM coverslips (**Fig. 2C**, right). At 300 pM TNF-α, the fractional contribution of background dAb signal is expected to be reduced due to the increased number of true binding events. As such, absolute dAb counts and colocalized and normalized counts yielded comparable CVs (**Fig. 2C**). Since combining colocalization and normalization produces a substantial increase in signal consistency in high-background conditions as well as across coverslips, we have used this analytical approach for the subsequent SiMCA analyses presented below.

### SiMCA lowers quantification errors in buffer and serum

Biological specimens such as serum and blood can generate especially high levels of non-specific background due to interferent proteins exhibiting cross-reactivity to dAbs or binding to the assay substrate itself. We evaluated how interfering species might impact the quantitative performance and precision of SiMCA versus a conventional single-color assay by comparing single-color and colocalization methods when generating calibration curves for 0–20 nM TNF-α in buffer and 70% chicken serum (**Fig. 3A**). We used chicken serum to mimic the complexity of human serum without interference from endogenous human TNF-α^12^.

**Figure 3:**
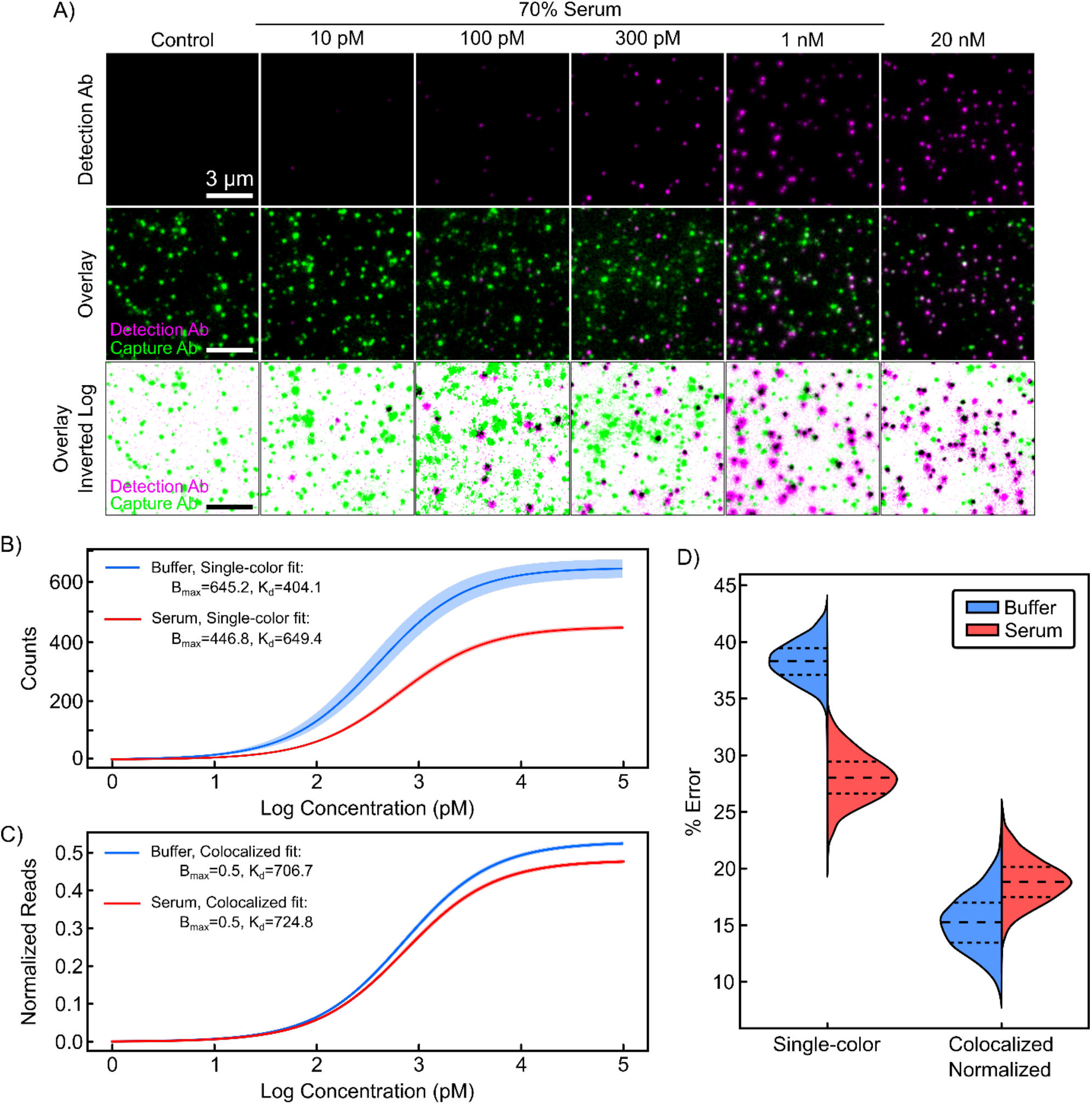
Quantification of TNF-α via single- and two-color analysis in buffer and 70% chicken serum. **A)** Single-color (*top*) and colocalized (*middle, bottom*) signal for increasing concentrations of TNF-α spiked into 70% chicken serum with 50 nM dAb. Best-fit TNF-α binding curves for **B)** absolute dAb counts and **C)** normalized, colocalized counts in buffer and 70% chicken serum. 2σ confidence curves are shaded. For primary data, please see **SI Fig. S7. D)** Bootstrapped error in TNF-α quantification for buffer and serum for absolute vs. normalized, colocalized dAb counts.

In an ELISA, quantification is achieved by fitting a Langmuir isotherm to data from samples spiked with known amounts of target using two fit parameters: an equilibrium dissociation constant (K_D_) and the maximum specific binding (B_max_). These parameters and their associated uncertainties are used to estimate the concentration of an unknown sample within a certain confidence interval. High confidence is gained when calibration curves have tight parameter fits that are unaffected by the sample matrix. Tighter parameter fits and lower background signal additionally lead to improved ability to resolve lower analyte concentrations, resulting in a lower LOD.

Using absolute dAb counts in buffer, we derived a K_D_ of 404 ± 35 pM and a B_max_ of 645 ± 16 counts (**Fig. 3B**). In serum, the same analysis yielded markedly different K_D_ and B_max_ values of 649 ± 17 pM and 446 ± 3.4 counts, respectively. The LOD also increased three-fold, from 6.6 ± 1.4 pM in buffer to 19.4 ± 4 pM in serum. Surprisingly, the CVs for these parameters were significantly lower in serum than in buffer. Consistent with observations from individual FOVs and coverslips, normalization to cAb counts rectifies this discrepancy, underlining the importance of coverslip-to-coverslip variation as a source of error if left uncorrected. Combining colocalization and normalization yielded narrower confidence envelopes as well as parameter fits for K_D_ and B_max_ that were virtually identical in serum and buffer (**Fig. 3C)**. In addition, we observed CVs that were 2.8–6.3-fold lower for the fitted parameters relative to values derived from absolute dAb counts, and these were roughly equivalent for serum and buffer experiments (see **SI Table 1)**. We also achieved a lower LOD in serum of 7.6 ± 2.0 pM, versus 19.4 ± 4 pM for the single-color measurement.

To estimate errors and the associated confidence in the quantification of unknown TNF-α concentrations, we implemented a bootstrapping approach using the above calibration data in buffer and serum (see Methods). Briefly, we used a subset of the data from each serum and buffer calibration as a training set, with the remaining data used as the test set. Using only the training dataset, we calculated new K_D_ and B_max_ parameter fits. We then used these fit parameters to predict sample concentrations for the test data set. The mean error was then calculated from the predicted versus true concentrations for the test set. We then repeated the process of splitting, fitting, testing, and calculating errors 1,000 times to obtain confidence intervals in the errors (see Methods). As shown in **Figure 3D**, normalized and colocalized dAb data reduced errors relative to single-color dAb counts by 2.5-fold and 1.5-fold for buffer and serum respectively (see **SI Table 2**). Notably, the errors with colocalization remained similar in buffer and serum.

### SiMCA lowers false-positive rates in complex samples

Finally, we set out to quantitively evaluate the diagnostic sensitivity and specificity of the single-color versus the two-color colocalization approach in a series of assays performed with low concentrations of TNF-α (0, 10, and 100 pM) spiked into either 70% chicken serum or 70% human blood. For reference, physiological concentrations of TNF-α range from 4 pM at baseline^13^ to a mean of 40 pM and up to 300 pM for patients developing septic shock^14^. In our experiment, absolute dAb counts varied widely in both serum and blood, with some distributions heavily skewed with long tails or even bimodal—this reflects the inherent heterogeneity of the coverslips, as discussed above. In contrast, normalized, colocalized counts were markedly more consistent across buffer, serum, and blood (see **SI, Fig. S8)**.

To evaluate the usefulness of colocalization and normalization in the context of a potential clinical application (*i*.*e*., detection of TNF-α detection), we conducted separate binary classifications between distributions with TNF-α and the control distribution without TNF-α. This enabled us to characterize the trade-off between true positive rates (TPR, or sensitivity) and false positive rates (FPR, or 1 – specificity) via receiver operating characteristic (ROC) curves. For reference, an ideal assay with little to no overlap between the target and control distributions would achieve high TPR with a low FPR – approaching perfect discrimination in the upper left corner (TPR = 1, FPR = 0). An assay with poor discrimination (*i*.*e*., producing similar distributions for both classes) would have an ROC curve closer to the diagonal (dotted line in **Figure 4A**), equivalent to random guessing.

**Figure 4:**
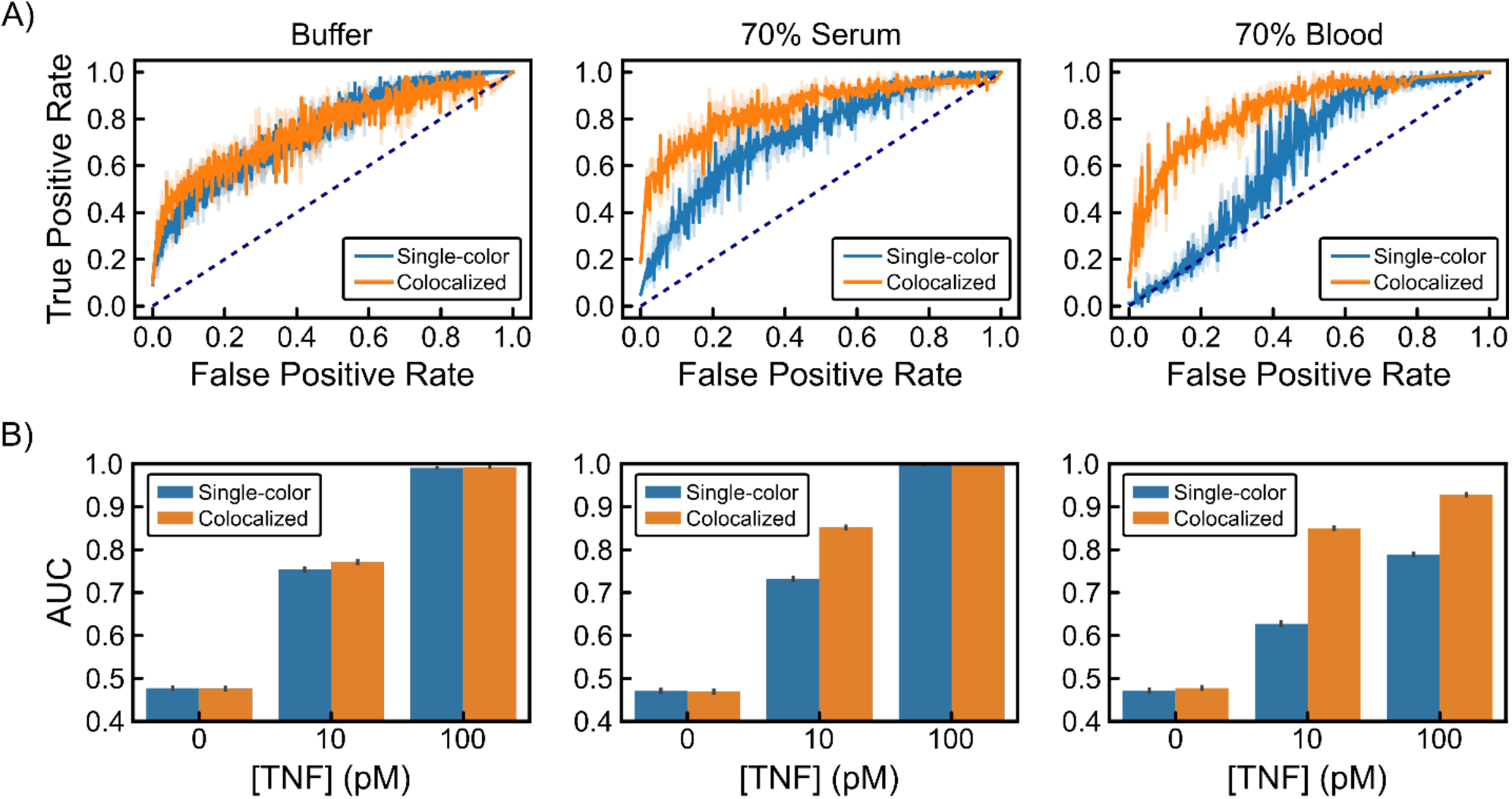
Discriminating TNF-α concentrations using single-color and colocalized methods in buffer, 70% chicken serum, and 70% human blood. **A)** Bootstrapped ROC curves from binary classification between 10 pM TNF-α distributions and TNF-α-free negative controls in buffer (*left*), 70% chicken serum (*middle*), and 70% human blood (*right*). Dotted lines indicate random guess. Raw data in **SI Figure S7. B)** Area under the ROC curve values from binary classification for 0, 10, and 100 pM TNF-α. Error bars are standard deviations from 1,000 bootstrapped samples. All distributions are from 128 FOVs collected from two coverslips per condition.

At 10 pM TNF-α, the ROC curves for the absolute dAb counts and colocalized, normalized methods were similar for samples prepared in buffer **(Fig. 4A)**. However, the colocalized, normalized data were clearly superior to those from absolute dAb counts in 70% serum or whole blood, particularly at lower FPRs, further highlighting the inability of single-color methods to distinguish false positives resulting from background dAb binding and coverslip variability **(Fig 4A; SI Fig. S8)**. In addition, the colocalized, normalized ROC curves were largely identical across buffer, serum, and whole blood **(Fig. 4A)**. To quantitatively compare the ROC curves across sample matrices and TNF-α concentrations, we calculated the area under the curve (AUC) (**Fig. 4B**). Increased TNF-α concentrations yielded an appreciable increase in AUC, as would be expected when distributions are being pushed further apart by increased numbers of true binding events. However, AUC values calculated from the single-color dAb counts were significantly lower than those derived from colocalized, normalized counts in both serum and blood. This difference is particularly apparent at 10 pM TNF-α, where nonspecific recruitment of dAb to the coverslip surface accounts for a considerable fraction of total counts. In summary, SiMCA achieved consistent and accurate analyte detection that was robust against background signal arising from complex biological specimens compared to a conventional single-color approach.

## Conclusion

In this work, we demonstrate that SiMCA can overcome the confounding effects of non-specific background to enable accurate, sensitive, and reproducible detection of picomolar protein concentrations even in highly complex sample matrices. We evaluated SiMCA using well-characterized, commercially available TNF-α antibodies, and found that the use of colocalization and normalization reduced variability and achieved a more consistent signal across coverslips, yielding CVs that were 2.8–6.3-fold lower than those derived based on absolute, single-color dAb counts. Our approach also produced quantification values that remained consistent in different sample matrices, with narrower confidence envelopes, more consistent parameter fits between serum and buffer, and a consistently lower LOD even in complex samples such as chicken serum and human blood. Thus, our technique provides a generalizable way to achieve far more robust immunoassay performance.

Our focus in this work was to understand and reduce general sources of error in immunoassays, rather than to demonstrate sensitivity that outperforms existing molecular detection assays. In theory, the sensitivity of SiMCA is limited only by the number of dAbs and cAbs counted, as well as the accuracy with which colocalization is determined. The former can be addressed simply by scanning larger areas of the coverslip—in the present study, we examined only ∼0.5% of the surface. Improvements to colocalization accuracy would help to eliminate false positives and allow higher cAb densities. As single-molecule fluorescence colocalization can be determined with Ångstrom precision^15^, the ultimate limit on colocalization in our assay would be the size of the antibodies used (∼10 nm). Förster resonance energy transfer (FRET) could also provide an alternate, stringent test of fluorophore colocalization, and we have noted the occurrence of FRET between colocalized dAbs and cAbs in the present study (**SI Fig. S9**). This mechanism could therefore offer an attractive target for future investigations. Finally, it should be noted that SiMCA requires relatively expensive microscopy equipment that can achieve single-molecule sensitivity. Extending the benefits of SiMCA to resource-limited environments will require strategies for boosting the fluorescence signal to levels that can be detected by smartphone cameras—for example, by using fluorescent nanoparticles that emit a substantially brighter signal, or fluorescence-enhancing materials that maximize the output from individual fluorophores.^16^ These would also be worthwhile avenues of investigation towards the larger goal of achieving ultra-sensitive, accurate, and reproducible molecular detection with readily available tools.

## Acknowledgements

This work was supported by the Chan-Zuckerberg Biohub, the Helmsley Trust, and the National Institutes of Health (NIH, OT2OD025342, R01GM129314-01).

## METHODS

### Materials and buffers

mAb1 (cAb) and mAb11 (dAb) anti-TNF-α antibodies were purchased from Biolegend. Human TNF-α protein was purchased from R&D Systems (210-TA). Huma blood was purchased from BioIVT. The SiteClick biotin antibody labeling kit, Alexa Fluor 546 NHS ester (succinimidyl ester), and all other chemicals were purchased from Thermo Fisher Scientific. All chemicals were of analytical grade and used without further purification. 1% v/v Vectabond was purchased from Vector Laboratories. PEG succinimidyl valerate MW-5000 (mPEG-SVA) and biotin-PEG-SVA MW-5000 (Biotin-PEG-SVA) were purchased from Laysan Bio. Custom imaging chamber components were purchased from Grace Bio-Labs. Coverslips were purchased from Thermo Fisher Scientific. The PBST buffer (pH 7.4) used in these experiments contained 137 mM NaCl, 2.7 mM KCl, 10 mM Na_2_HPO_4_, 1.44 mM KH_2_PO_4_, and 0.1% Tween 20. The 1X PBSBT buffer contained 137 mM NaCl, 2.7 mM KCl, 10 mM Na_2_HPO_4_, 1.44 mM KH_2_PO_4_, 0.1% Tween 20, and 1% BSA.

### Sample preparation and imaging

Coverslips were soaked in piranha solution (25% H_2_O_2_ and 75% concentrated H_2_SO_4_) and sonicated for 1 h, followed by multiple rinses in water (Thermo Fisher Scientific, molecular-biology grade) and acetone (Thermo Fisher Scientific, HPLC grade). Dry and clean coverslips were then treated with Vectabond/acetone (1% v/v) (Vector Labs) solution for 5 min and then rinsed with water and left in a dried state until used. In order to prevent non-specific adsorption of biomolecules onto the glass surface, coverslips were functionalized prior to use with a mixture of poly(ethylene glycol) succinimidyl valerate, MW 5,000 (mPEG-SVA) and biotin-PEG-SVA at a ratio of 99:1 (w/w) (Laysan Bio) in 0.1 M sodium bicarbonate (Thermo Fisher Scientific) for 3 h.^10^ Excess PEG was rinsed with water, and the coverslips were dried under a N_2_ stream. Imaging chambers (∼5 μL) were constructed by pressing a polycarbonate film (Grace Bio-Labs) with an adhesive gasket onto a PEG-coated coverslip. Two silicone connectors glued onto the pre-drilled holes of the film served as inlet and outlet ports. The surface was incubated with 7 μL of a 2 mg/ml neutravidin solution (Thermo Fisher Scientific). Excess neutravidin was then washed off with 100 μL of 1X PBS buffer.

### TIRF microscopy

Single-molecule fluorescence measurements were performed with objective-type TIRF microscopy on an inverted microscope (Nikon TiE) with an Apo TIRF 100x oil objective lens, NA 1.49 (Nikon) as described previously^17^, and controlled using Micro-Manager^18^. Samples were excited with a 532-nm (Crystalaser) or 635-nm (Blue Sky Research) laser. Excitation light was cleaned with a quad-edge laser-flat dichroic with center/bandwidths of 405/60 nm, 488/100 nm, 532/100 nm, and 635/100 nm from Semrock (Di01-R405/488/532/635-25×36), and the emission signal was passed through the corresponding quad-pass filter with center/bandwidths of 446/37 nm, 510/20 nm, 581/70 nm, 703/88 nm (FF01-446/510/581/703-25). The emission signal was then separated using a dichroic beam splitter (635 nm), passed through an additional set of filters (546 channel: 593 nm/40 nm (Semrock); 647 channel: 675/30 nm (Semrock), and recorded on an EMCCD camera (Andor iXon), as described previously^17^. We were capturing 16-bit 512 × 512 pixel images with an exposure time of 200 ms, and a multiplication gain of 2800-3000. Excitation was carried out at a full power setting (25 mW) with a power output of 2-3.5 mW at the objective for the green (532 nm) laser. The excitation power of the red laser ranged between 2 and 4 mW at the objective based on the experiment. We typically observed 150–300 spots per 35 µm x 70 µm field of view.

### Antibodies

The anti-human TNF-α cAb (mAb1) was functionalized in house. The antibody was first biotinylated site-specifically using the SiteClick biotin antibody labelling kit according to the manufacturer’s instructions. The SiteClick biotin antibody labeling kit specifically attached the biotin to the heavy chain of the antibody, targeting the carbohydrate domains present on essentially all IgG antibodies and thereby ensuring that the antigen-binding domains remain available for binding to the antigen target. The antibody was then labeled with the Alex Fluor 546 antibody labeling kit (Thermo Fisher Scientific, A20183) according to the manufacturer’s instructions **(SI Fig. S1)**. We typically observed 2.5–3 Alexa 546 fluorophores per antibody. Anti-human TNF-α dAbs (mAb11) were pre-labelled with Alexa Fluor 647 and used at the indicated concentrations. Quality control experiments confirmed that 1) binding affinity of antibodies was not compromised by labelling, 2) labelling efficiency of the different reagents is close to 90%, 3) non-specific background is caused mainly by dAbs and cAbs adhering to the surface, with minimal TNF-α sticking to surface even at very high target concentrations.

### Choice of fluorescent labels

We chose Alexa 546 for this assay because it has greatly improved photostability over Alexa 532 under our experimental conditions, where rapid photobleaching and/or blinking could compromise data analysis and event counting. We found that the average survival time for Alexa 546 was notably larger than that of Alexa 532—14 s versus 3 s—in the assay conditions that we chose to minimize blinking. We thus concluded that Alexa 546-labeled cAbs will not photobleach within a time-frame that would interfere with the counting process **(SI Fig. S10)**.

### Detection of TNF-α

Neutravidin-coated coverslips were first incubated with 7 μL of 0.3 nM biotinylated Alexa 546-labeled cAb solution for five minutes, then washed using 1X PBST buffer to get rid of unbound material. Images of coverslips before and after cAb addition were acquired to account for background noise, channel leakage, and donor fluorophore intensity (see **SI Fig. S9)**. Subsequently, we added 7.5 μL of 50 nM dAb solution spiked with recombinant human TNF-α at different concentrations (0, 0.01, 0.1, 0.3, 1, and 20 nM). The TNF-α was prepared as 25 µl samples in 1.5 mL Protein LoBind tubes (Eppendorf). TNF-α was thawed on ice and added at 5.9x (for 0.01, 0.1, 0.3, and 1 nM) or 2.95x (for 20 nM) the final concentration. PBSBT was added to a final volume 17.5 µl, after which 7.5 µl of dAb was added for a final dAb concentration of 50 nM. The samples were immediately injected into the prepared coverslips and incubated covered overnight at 4 °C. For chicken serum and human blood experiments, we replaced the buffer in the above protocol with chicken serum or human blood, where the final serum and blood percentage after mixing with dAb was 70%. The fluorescent background greatly increased in the presence of serum or blood in the coverslip due to autofluorescence/quenching effects, but an additional washing step restored the initial signal/background ratio. **SI Figure S11** shows that the distribution of cAb intensities and counts remained constant following overnight incubation with buffer and serum. This demonstrates the robustness of the surface passivation layer, avidin linkage, and fluorophore-antibody complex.

We note that because TNFα is a small protein, we could detect Förster resonance energy transfer (FRET) between the donor fluorophore on the cAb and acceptor fluorophore on the dAb upon binding the protein target. Because we were working with Alexa 546 and Alexa 647, with an R0 = 8 nm, we were able to detect energy transfer and compute FRET efficiencies upon protein binding. This was confirmed by the drop in donor intensity and increase in acceptor intensity upon target binding **(SI Fig. S9)**. The FRET signal was discarded as it is beyond the scope of this study.

### Image segmentation and registration

We stitched together an image to directly map cAbs and dAbs from the green- and red-only excitation images (green left, red right). We denoised by performing a gaussian filter using a standard deviation for the kernel of 0.8. We then isolated regional maxima. Since the raw images can be unevenly illuminated, we first subtracted from the gaussian-filtered image a background image obtained by morphological reconstruction, as described in the *scikit-image* example^19^. The resulting background subtracted image was then mapped to a more dynamic intensity range by an inverse hyperbolic sine transformation. Local maxima and their respective x, y coordinates were selected if their intensity values were at least 1.2 times the standard deviation plus the median. cAb and dAb spots were differentiated by locations of the detected local maxima in the left and right halves of the stitched image, respectively. To determine the colocalized spots, we mapped the coordinates of the dAb spots to the region of the cAb using a pre-defined affine transformation (see below). The transformed dAb coordinates (dAb’) were then matched to the true detected cAb coordinates. If a pair of coordinates (cAb[i], dAb’[j]) were within a Euclidean distance of 1.5 pixels, the pair was counted as a colocalized spot.

The affine transformation matrix was made per experiment day to account for any misalignment of the microscope setup. First, 100-nm Tetraspeck beads (Thermo Fisher Scientific T7279) were rastered across a glass coverslip. Five to ten images were taken, and a transformation matrix was determined for each image and averaged to produce the final transformation matrix. Capture and detection images were stitched and localized as described above. Capture and detection coordinates were matched up to their closest neighbors across all coordinates using a KD tree with leaf size of 30 and using the Euclidean distance metric. Coordinates in both images were saved as C and D matrices, respectively, of size *n* x 3 where *n* is the number of beads detected. The coordinates c_0_, …, c_n_ were projected onto the corresponding coordinates d_0_, …, d_n_ using the affine transformation matrix T:

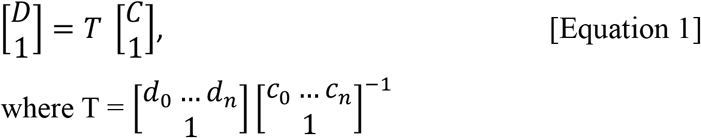

The average of T matrices was calculated for each of the bead images and provided for transformation of experimental data.

Finally, we applied a quality filtering step. If the spots were not well-distributed, this could reflect faulty frames of view due to bubbles, large dust particles, etc. Since the dynamics of target and dAb binding to the coverslip surface could be significantly altered in these frames, we removed these in this post-processing step. The mean *y* coordinates of the spots detected in the left half of the image were averaged. If the averaged coordinate was not within 75 pixels of the middle of the image, that frame of view was removed from analysis. Less than 1% of frames of view were discarded because of this filter.

### Absolute single-color and normalized, colocalized analysis

Absolute single-color dAb spots were simply localized and counted in the Alexa-647 channel. However, this counting is highly dependent on the consistency of cAb coverage across coverslips and frames of view. For SiMCA colocalized spot analysis, we addressed coverslip variability by normalizing spot counts relative to the cAb signal in the Alexa-546 channel:

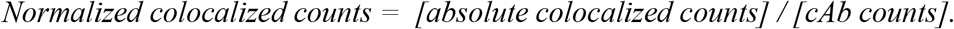

For comparison of just normalization methods to absolute single-color dAb spots in our SI, we also normalized single-color counts as such:

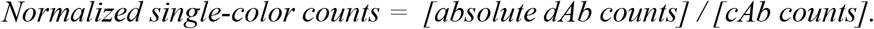

We found normalization lowers quantification error, and thereby enable us to exploit colocalization to reduce the effects of non-specific binding.

### Fitting to binding curve

To create a calibration curve, we fitted our data to the Langmuir binding isotherm:

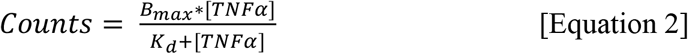

where counts can be single-color unnormalized counts, normalized two-color counts, or normalized, colocalized counts, and [*TNFα*] is the concentrations used in the binding curves (10 pM, 100 pM, 300 pM, 1 nM, and 20 nM). K_d_ (equilibrium dissociation constant) and B_max_ (maximum signal possible) were determined by the curve-fitting function in python using Scipy’s ‘optimize curve fit’ function, which uses non-linear least squares to fit a function. To calculate the concentration given a signal, we then used the inverse function with fit parameters K_d_ and B_max_ as follows:

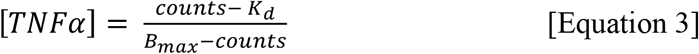

### Calculation of LOD

As per convention, we defined LOD as the signal that is three standard deviations (*σ*_*y*_) above the mean signal 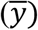 obtained without analyte: 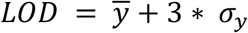. We then converted this value to its associated concentration using the fits from the binding curve and the inverse binding curve function (**Eq. 3**). The corresponding error in LOD is determined by propagation of errors of the inverse function (*f*) and errors associated with the binding curve fits:

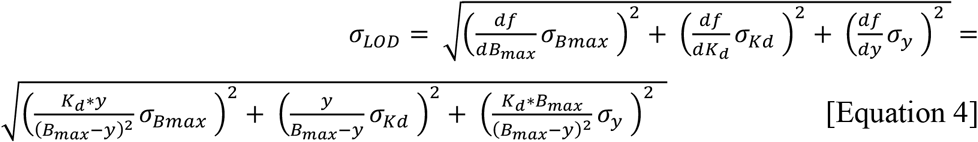

### Quantification with Bootstrapping

We bootstrapped our data 1,000 times to properly characterize mean errors and confidence in quantification of new unknown targets. For each bootstrap iteration, we sampled with replacement from our dataset of 128 FOVs, fit to the binding curve in **Equation 2**, then used the remaining unsampled data to predict the TNF-α concentration for each FOV (**Eq. 3**). We calculated the error between the predicted and true concentration of these test samples using Mean-Absolute-Percentage-Log-Error (MAPLE):

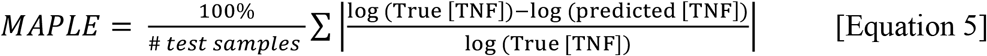

Finally, after all bootstrap iterations, we calculated the standard deviation of MAPLE.

### Bootstrapped binary classification, ROC, and AUC calculations

We conducted binary classification given two distributions to quantitively evaluate diagnostic sensitivity and specificity. First, we combined the two distributions into one dataset with N frames of view: {(x_1_, y_1_), …, (x_N_, y_N_)} (typically N =128). A given single-color or colocalized count, x_i_, is assigned to a label, y_i_, where y_i_ is 0 if it is the 0 pM control or 1 if it is a sample with TNF-α. We then split the dataset into training and test datasets, T_train_ and T_test_. Using the sklearn python library^20^, we fitted our training data to a binary logistic regression classifier without regularization, such that we minimize the following cost function with respect to the parameters *w* and *c*:

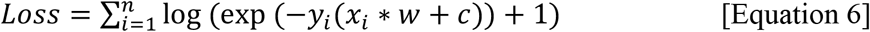

Using the T_test_ dataset and sklearn libraries, we then calculated the probability estimates and respective ROC curve and AUC values. To estimate confidence in the ROC and AUC values, we bootstrapped the above binary classification 1,000 times. For each bootstrap iteration, we sampled each distribution with replacement for the training set. The remaining unsampled data were then used as the test set. The above classification and ROC/AUC metrics were then calculated.

## Notes

### Competing Interest Statement

The authors have declared no competing interest.

