## Supporting information for "Improved immunoassay sensitivity and specificity using single-molecule colocalization"

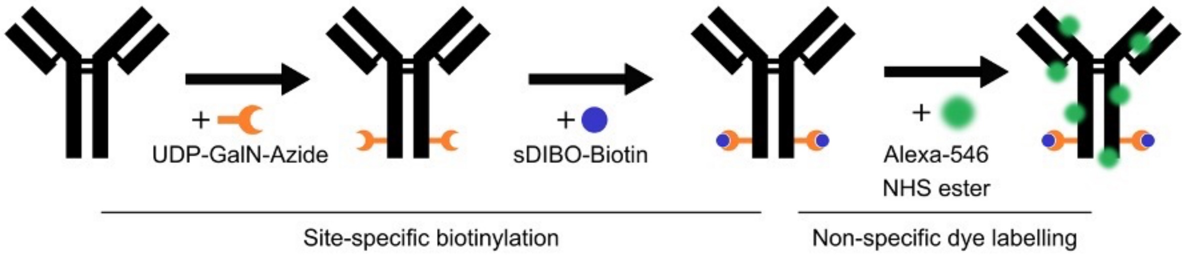


**Figure S1:** Strategy for site-specific and non-specific cAb labeling with fluorophores and biotin labels.

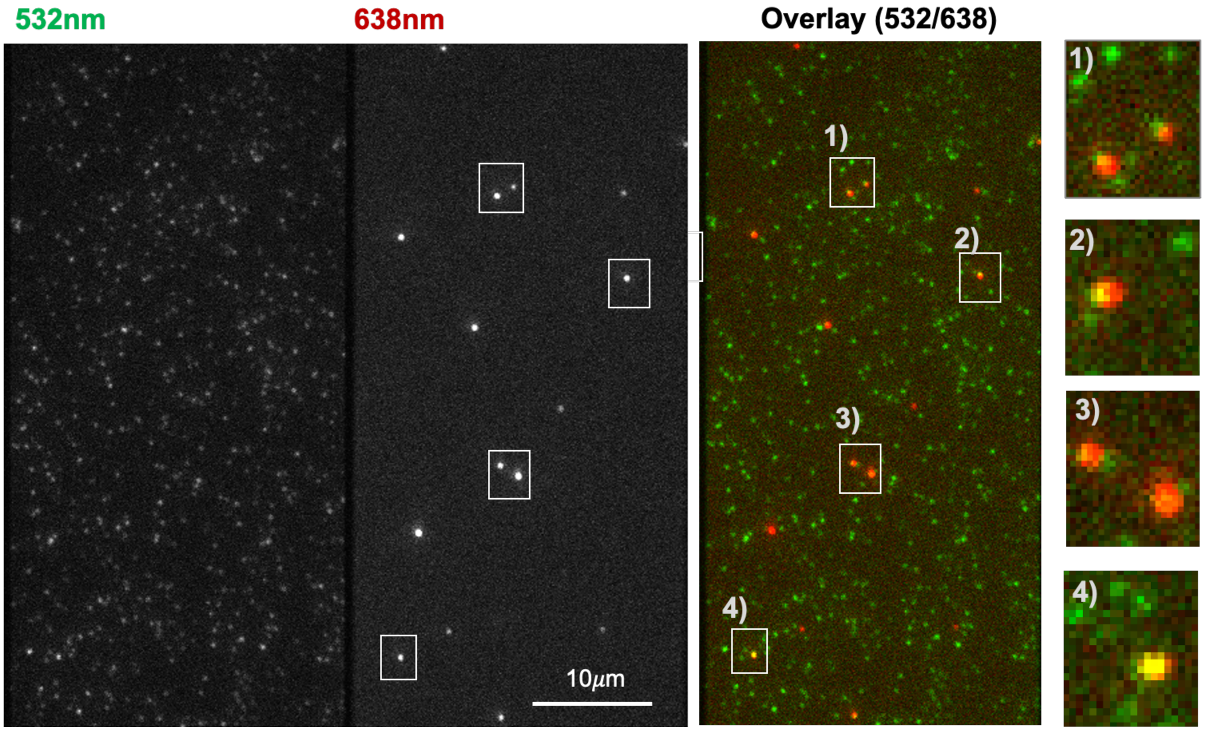


**Figure S2:** Fluorescence images of Alexa 546 (left), Alexa 647 (middle) and the overlay of those two channels (right) showing specific binding at low concentrations (50 nM) of dAb in the presence of 100 pM TNF-α in buffer. Far-right panels show magnification of labeled regions of interest.

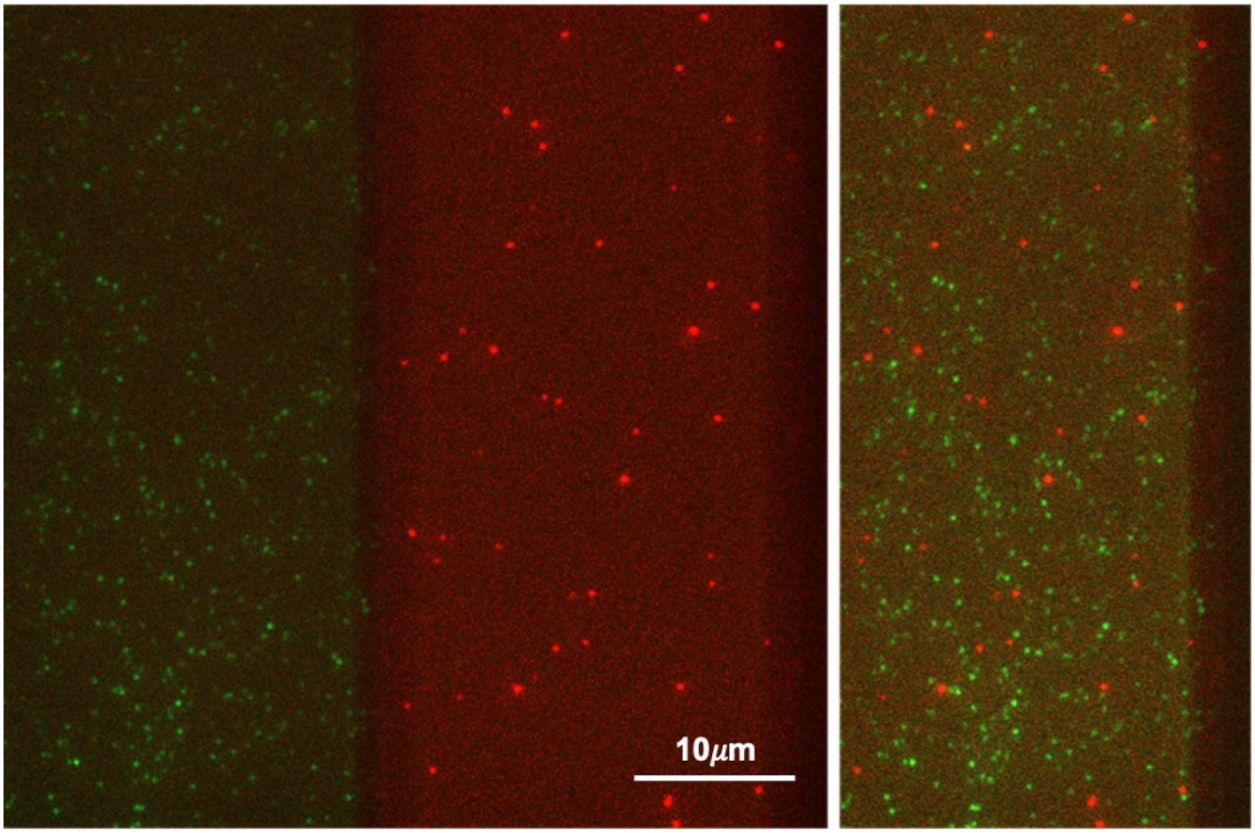


**Figure S3:** Fluorescence images of Alexa 546 (green, left), Alexa 647 (red, middle), and overlays of the two channels (right) showing high levels of non-specific binding at high concentrations (800 nM) of dAb in the absence of TNF-α**.**

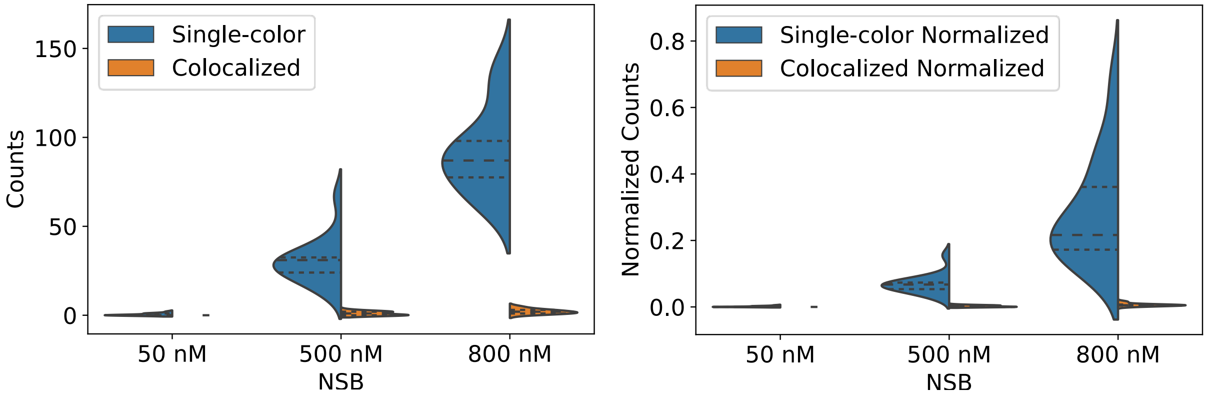


**Figure S4:** Number of counts observed with increasing concentrations of dAb (50, 500, and 800 nM) in the absence of TNF-α using single-color and two-color analysis approaches with absolute counts (left) and cAb-normalized counts (right).

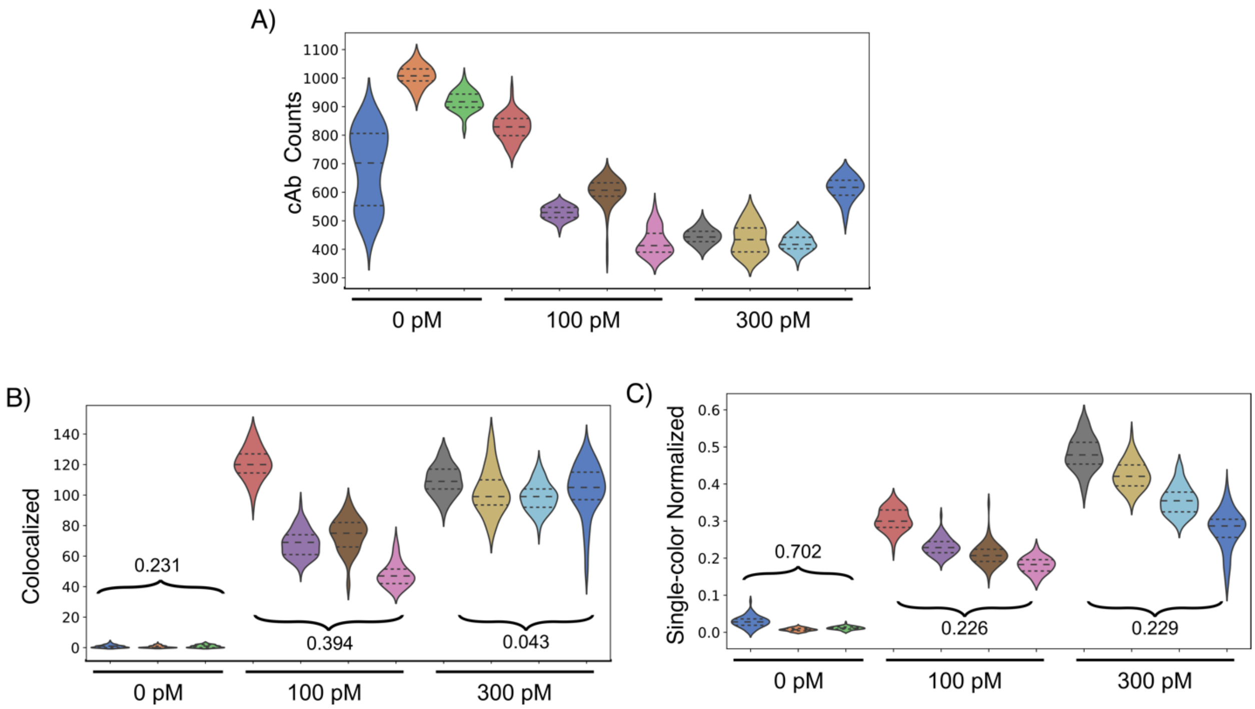


**Figure S5:** A) Distribution of absolute cAb counts, B) colocalized dAb and cAb counts, and C) single-color normalized dAb counts across 11 coverslips and three different TNF-α concentrations. Numbers in plots B and C present CVs for a given condition.

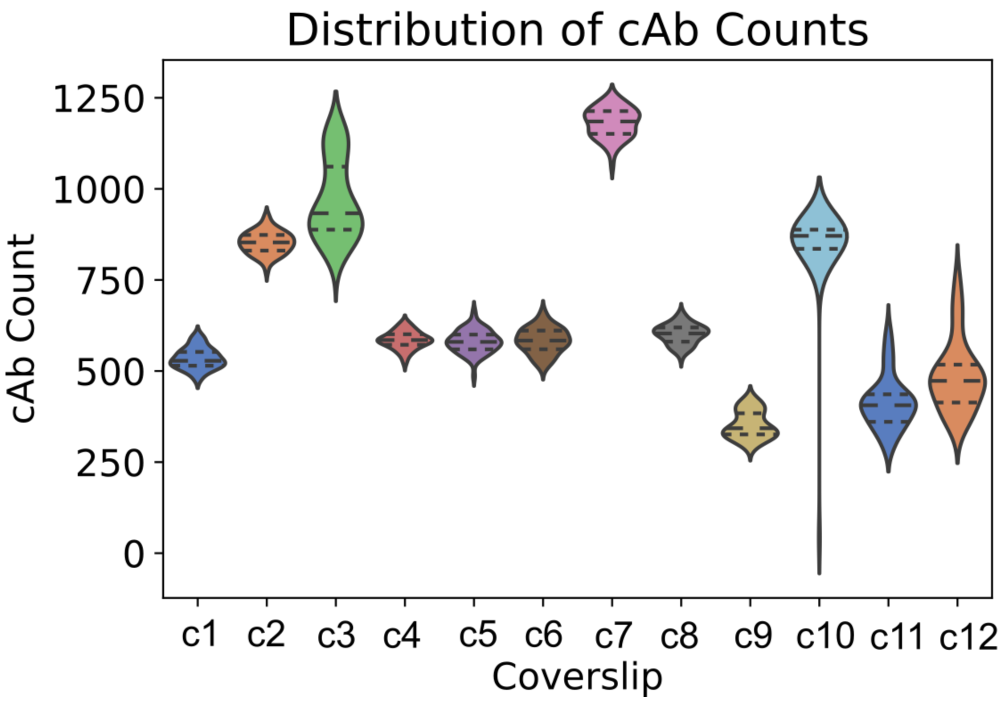


**Figure S6:** Heterogeneity in the distributions of Alexa Fluor 546-labeled cAb. Number of counts across 12 different coverslips (128 different FOVs from each coverslip) after an overnight incubation with target/dAb in buffer.

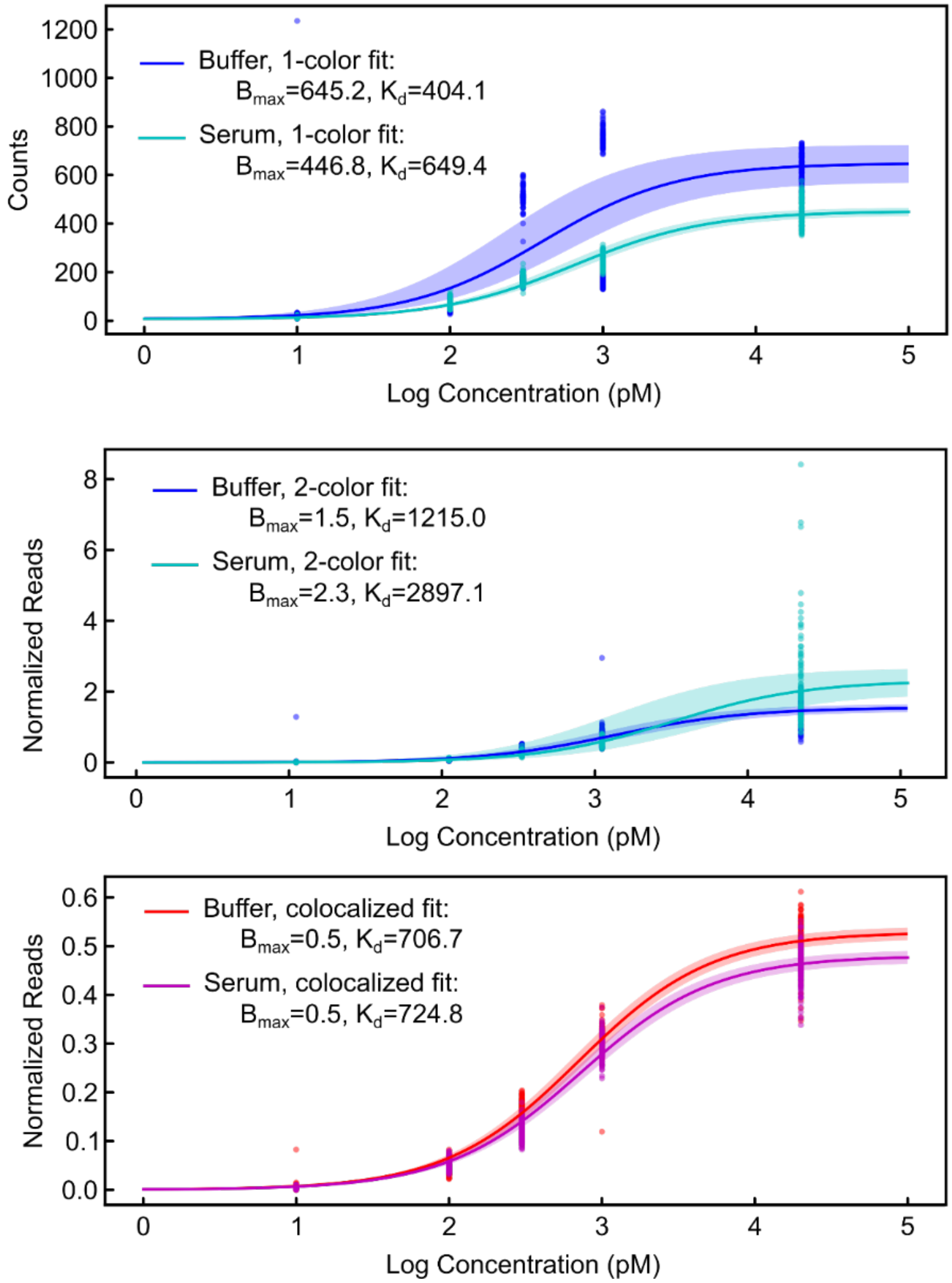


**Figure S7:** Absolute single-color dAb, single-color normalized, and colocalized, normalized counts from each FOV and their respective binding curve fits. 2-sigma confidence curves are shaded.

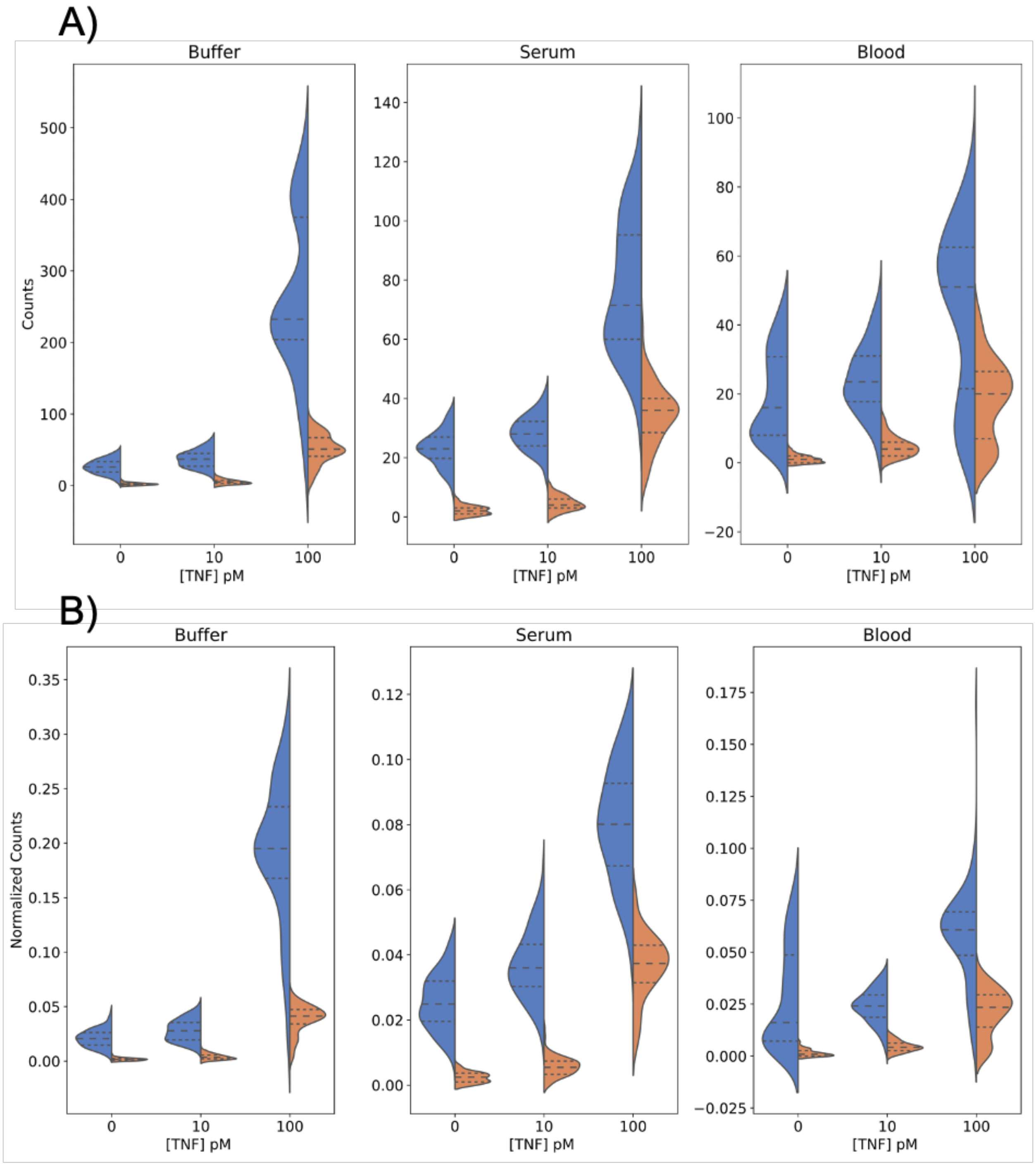


**Figure S8:** Distribution of (A) absolute and (B) normalized single-color dAb (blue) and colocalized counts (orange) from 0, 10, and 100 pM TNF-α in buffer (left), 70% chicken serum (middle), and 70% blood (right). Dashed lines separate quartiles of the distribution.

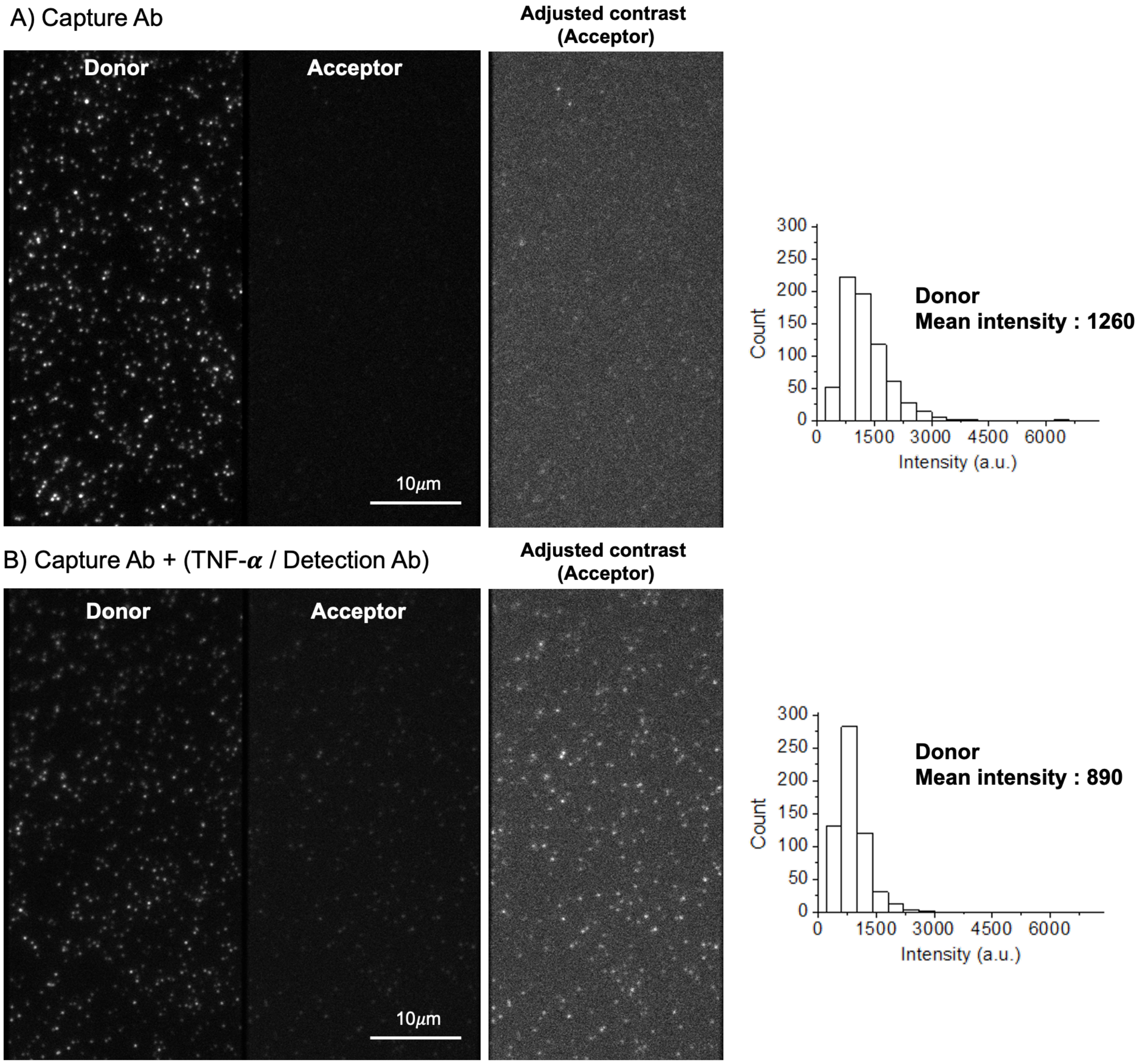


**Figure S9:** Because TNF-α is a small protein, we can detect Förster resonance energy transfer (FRET) between donor fluorophores on the cAb and acceptor fluorophores on the dAb upon binding the protein target. A) Fluorescence images (left) and donor mean intensities (right) of cAb upon 532 nm excitation before and B) after target/dAb incubation. The average intensity dropped from 1,260 to 890 after adding target/dAb, and we also observed an increase in the intensity of the acceptor channel upon 532 nm excitation. Adjusted contrast images for the acceptor channel are shown on the right to highlight the intensity increase in the acceptor channel. Scale bar = 10 μm.

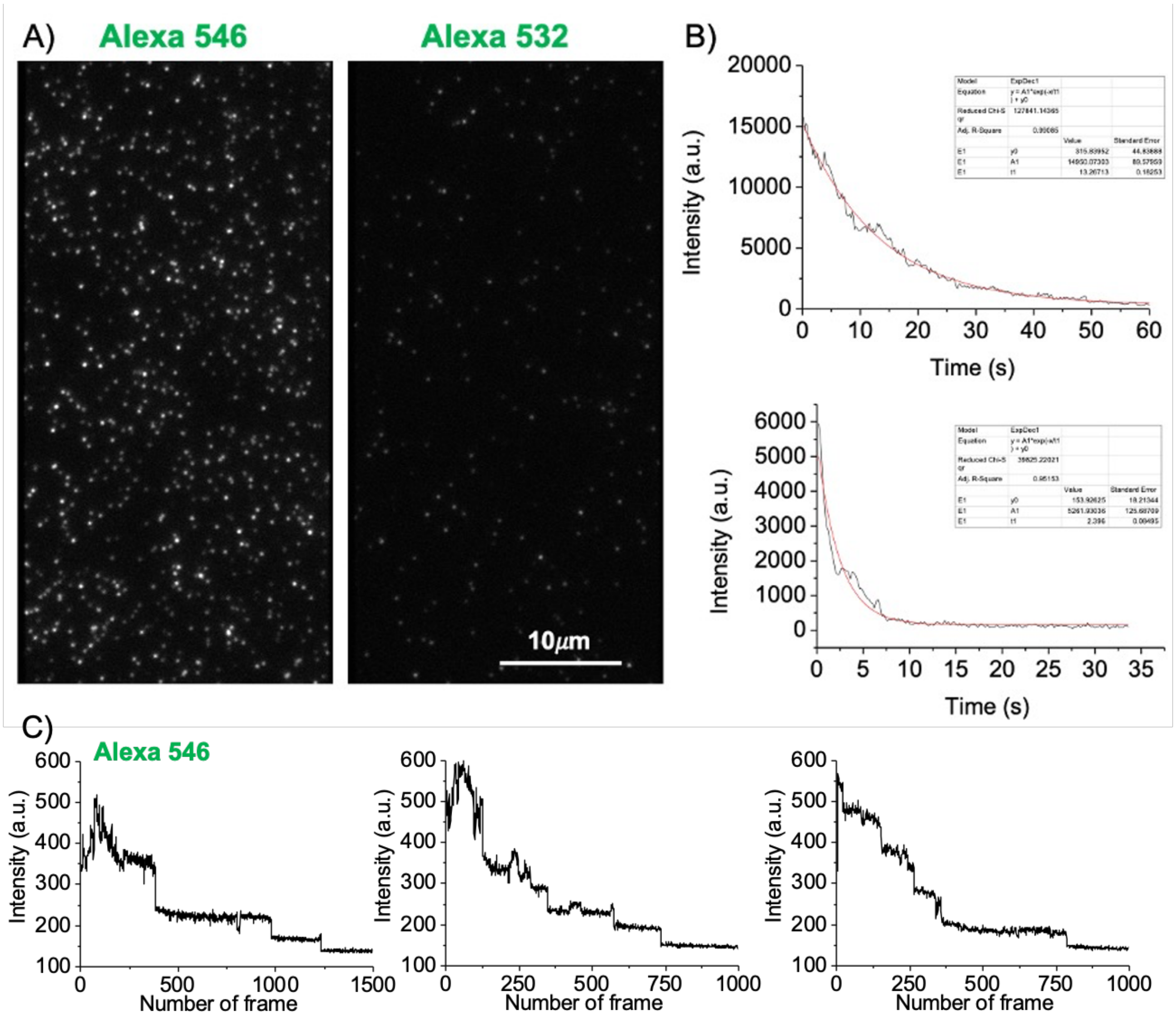


**Figure S10:** Lifetime analysis of Alexa 546- and 532-labeled cAbs with TIRF microscopy. (A) Single image (raw data) of Alexa 546 and Alexa 532 spots on the coverslip surface after excitation at 532 nm for 2 min (at exposure = 200 ms, gain = 3,000, power = 3 mW; see Methods for details). Scale bar = 10 µm. (B) Photobleaching time recorded for Alexa 546 (top) and Alexa 532 (bottom). C) Examples of the intensity-time trajectories of bleaching of fluorescence emission for Alexa 546-labeled antibodies.

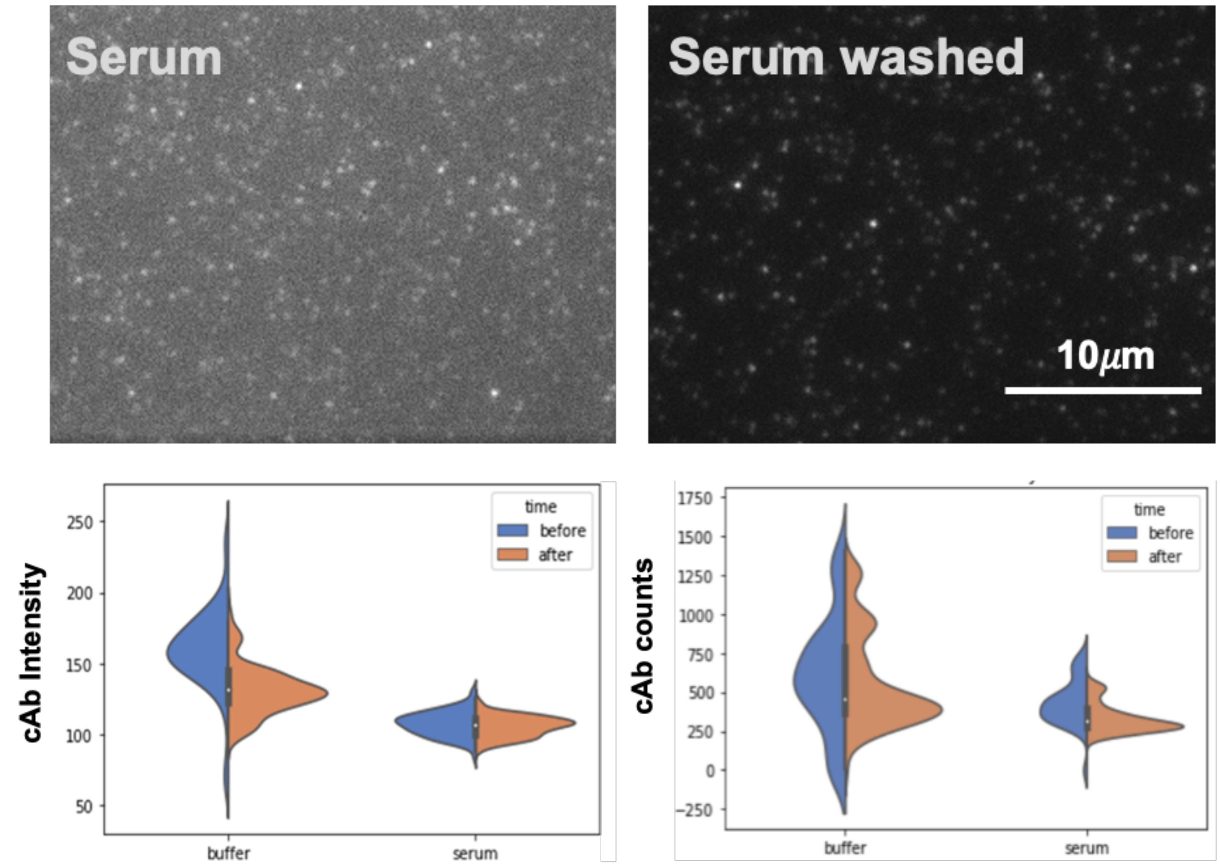


**Figure S11:** cAb fluorescence images in serum before (top left) and after washing with buffer (top right). Distributions of Alexa Fluor 546-labeled cAb intensity (bottom left) and number of counts (bottom right) across six coverslips before and after overnight incubation with buffer and serum. The distribution of cAb intensities and counts remained constant after overnight incubation with buffer and serum, which demonstrates the robustness of the system.

**Table 1:** Parameters and respective standard deviation errors from fitting to Langmuir isotherm using single-color and colocalized counts for both normalized and non-normalized methods.

| | Buffer | | | Serum | | |
|---|---|---|---|---|---|---|
| | Bmax (CV) | Kd (CV) | LOD (CV) | Bmax (CV) | Kd (CV) | LOD (CV) |
| **1-Color** | 645 ± 16 (0.024) | 404 ± 35 (0.088) | 6.6 ± 1.4 (0.210) | 446 ± 3.4 (0.008) | 649 ± 17 (0.026) | 19.4 ± 4 (.212) |
| **Colocalized** | 235 ± 4.1 (.017) | 288 ± 19 (.066) | 2.1 ± 0.6 (0.286) | 133 ± 2.1 (0.016) | 201 ± 12 (.060) | 4.9 ± 1.3 (0.256) |
| **1-Color norm** | 1.55 ± 0.02 (0.014) | 1,214 ± 59 (0.049) | 11.1 ± 2.2 (0.201) | 2.309 ± 0.07 (0.031) | 2,897 ± 347 (0.120) | 26.4 ± 5.8 (.219) |
| **Colocalized norm** | 0.528 ± 0.003 (0.005) | 706 ± 11.940 (0.017) | 3.1 ± 0.9 (0.290) | 0.480 ± 0.003 (0.005) | 725 ± 13.5 (0.019) | 7.7 ± 2.0 (0.258) |

**Table 2:** MAPLE Error from bootstrapping using single-color and colocalized counts for normalized and non-normalized methods. CV is displayed in parentheses.

| | Buffer | Serum |
|---|---|---|
| **1-Color** | 38 ± 1.7 (0.045) | 28 ± 2.1 (0.076) |
| **Colocalized** | 26 ± 2.4 (0.091) | 25 ± 1.9 (0.076) |
| **1-Color norm** | 28 ± 2.0 (0.071) | 29 ± 2.2 (0.078) |
| **Colocalized norm** | 15 ± 2.4 (0.160) | 19 ± 2.0 (0.105) |
